# Multi-Tissue Neocortical Transcriptome-Wide Association Study Implicates 8 Genes Across 6 Genomic Loci in Alzheimer’s Disease

**DOI:** 10.1101/2020.06.29.178590

**Authors:** Jake Gockley, Kelsey S. Montgomery, William L. Poehlman, Jesse C. Wiley, Yue Liu, Ekaterina Gerasimov, Anna K Greenwood, Solveig K. Sieberts, Aliza P. Wingo, Thomas S. Wingo, Lara M. Mangravite, Benjamin A. Logsdon

**Author notes:** Corresponding authors (B.A.L).

## Abstract

**Background:** Alzheimer’s disease (AD), an incurable neurodegenerative disease, currently affecting 1.75% of the United States population, with projected growth to 3.46% by 2050. Identifying common genetic variants driving differences in transcript expression that confer AD-risk is necessary to elucidate AD mechanism and develop therapeutic interventions. We modify the FUSION Transcriptome Wide Association Study (TWAS) pipeline to ingest expression from multiple neocortical regions, provide a set of 6780 gene weights which are abstracatable across the neocortex, and leverage these to find 8 genes from six loci with associated AD risk validated through summary mendelian randomization (SMR) utilizing IGAP summary statistics.

**Method:** A combined dataset of 2003 genotypes clustered to Central European (CEU) ancestry was used to construct a training set of 790 genotypes paired to 888 RNASeq profiles across 6 Neo-cortical tissues (TCX=248, FP=50, IFG=41, STG=34, PHG=34, DLPFC=461). Following within-tissue normalization and covariate adjustment, predictive weights to impute expression components based on a gene’s surrounding *cis*-variants were trained. The FUSION pipeline was modified to support input of pre-scaled expression values and provide support for cross validation with a repeated measure design arising from the presence of multiple transcriptome samples from the same individual across different tissues.

**Results:** *Cis*-variant architecture alone was informative to train weights and impute expression for 6780 (49.67%) autosomal genes, the majority of which significantly correlated with gene expression; FDR < 5%: N=6775 (99.92%), Bonferroni: N=6716 (99.06%). Validation of weights in 515 matched genotype to RNASeq profiles from the CommonMind Consortium (CMC) was (72.14%) in DLPFC profiles. Association of imputed expression components from all 2003 genotype profiles yielded 8 genes significantly associated with AD (FDR < 0.05); APOC1, EED, CD2AP, CEACAM19, CLPTM1, MTCH2, TREM2, KNOP1.

**Conclusion:** We provide evidence of cis-genetic variation conferring AD risk through 8 genes across six distinct genomic loci. Moreover, we provide expression weights for 6780 genes as a valuable resource to the community, which can be abstracted across the neocortex and a wide range of neuronal phenotypes.

## Intro

Alzheimer’s Disease (AD) is a progressive, incurable neurodegenerative disease accounting for 60-70% of all dementia diagnoses[1], currently affecting 5.8 million Americans and projected to grow to 13.8 million diagnoses by 2050[2]. Late Onset Alzheimer’s Disease (LOAD) comprises over 95% of AD diagnoses, and is composed of a diverse, largely unknown set of etiologies[3]. The mechanisms of AD remain insufficiently explained, despite clear A**β** plaque and Tau neurofibrillary tangle neuropathology. Additionally, known highly penetrant genetic variants from familial-based cohorts with Early-Onset Alzheimer’s Disease (EOAD) implicate genes such as APP, PSEN1, PSEN2. Despite clear pathology and known risk factors, AD therapeutic clinical trials have consistently failed[4,5]. Elucidating the mechanisms by which AD genetic risk loci contribute to AD disease and disease progression is instrumental in the development of future impactful therapeutics.

While genome wide association studies (GWAS) identified some candidate loci associated with AD risk, genes targeted through *cis-*genetic risk factors remain unclear [6,7]. Likewise, postmortem bulk-cell transcriptomics show vast expression changes across multiple neocortical regions, however it remains difficult to identify which differentially expressed genes are driving AD from gene expression changes caused by the disease state of widespread cell death and tissue degeneration[8,9]. Transcription-Wide Association Studies (TWAS) help provide this associative bridge and mechanistic direction of effect between genotype, transcript, and disease status[10]. TWAS leverages the cis-genotype region surronding a expressed gene to predict the *cis*-heritable component of a gene’s expression, which in turn can be associated to disease status using GWAS summary statistics.

We modified the FUSION pipeline[10], which deploys blup, bslmm, lasso, top1, and enet models to predict gene expression from *cis*-variants within 1MB of a given gene to train weights and impute expression for 6780 (49.67%) autosomal genes from matched genotypes and RNA-Seq profiles from Dorsolateral Prefrontal Cortex (DLPFC), Temporal Cortex (TCX), Prefrontal Cortex (PFC), Superior Temporal Cortex (STG), Inferior Temporal Gyrus (IFG), and Parahippocampal Gyrus (PHG) provided by the Accelerating Medicines Partnership - Alzheimer’s Disease (AMP-AD) consortia. Imputed gene expression validated in CommonMind Consortium (CMC) DLPFC. Using our trained models, we then imputed gene expression into a large, recent GWAS cohort[6] to identify genes showing differential predicted gene expression between AD patients and controls. Following correction multiple testing, Joint Conditional Probability testing (JCP), and Summary Mendelian Randomization (SMR) we discover eight candidate AD risk genes APOC, EED, CD2AP, CEACAM19, CLPTM1, MTCH2, TREM2, and KNOP1.

Expanding associated genes into gene sets using co-expression yielded enrichments for specific cell-type marker sets particularly microglial, oligodendrocyte, and astrocyte cell populations and cellular functions such as protease binding, myeloid and leukocyte regulation/activation, regulation lipid/lipoprotein, RNA splicing, and steroid regulation. We identify 8 genes across six distinct genomic loci associated to AD through gene expression attributable to their *cis*-genetic variation. Trained gene expression weights are a community resource which can be abstracted to multiple phenotypes, and gain further insights from large genotyped cohorts to maximize the informativeness of invaluable and rare patient material[11]. To this end, we provide a valuable resource to the community in the form of predictive gene expression weights which can be leveraged across a wide range of neurological phenotypes.

## Methods

### Ancestry Analysis

Ancestry analysis and clustering was performed to define CEU ancestral individuals from the combined ROSMAP, Mayo and MSBB cohort. Briefly, phaseI 1000 Genomes data[12] was filtered for YRI, CHB, JPT, and CEU ancestral populations. Genotype data was combined across MSBB, Mayo, and ROSMAP cohorts, filtered for 1000G overlapping SNPs, and combined with 1000G data from the four reference ancestral populations. PCA was performed using Plink (v1.9) on SNPs passing filtering; Minor Allele Frequency > 1%, Missingness < 0.1, maximum minor allele frequency < 40%, and independent pairwise linkage filter window of 50Kb at 5Kb steps and a r-squared threshold of 0.2. PCA results were visualized along PC1 (50.7%) and PC2 (31.6%). Genotype clustering to identify clustered genotype profiles was performed with the R package GemTools[13,14]. The clustering of genotype profiles was performed on all PCs describing greater than 1% of variation. Genotypes were recoded for GemTools clustering in plink similar to the PCA but with the added flags; --recode12, --compound-genotypes, --geno 0.0000001. Ancestrally matched CEU samples were identified as any sample genotypes belonging to one of the three out of eight clusters to which contained 1000G reference CEU individuals (Fig. S1).

### Defining The TWAS Training Set

Genotype data for 2360 individuals was combined across MSBB (N=349), Mayo (N=303), and ROSMAP (N=2360) cohorts *(Specific details see: supplementary methods sections 1-2)*. CEU 1000G reference individuals were present within the top 3 clusters (Fig. S1c) which contained 2003 individuals across all cohorts (Fig. S1[b,d]). Broader admixture was observed in the MSBB cohort compared to ROSMAP and Mayo representative of study recruitment and population (Fig. S1[e-h]). After filtering for variants present within the LD-Reference panel the final European ancestry filtered population of 2003 individuals were represented by 1,069,623 variants (Fig. 1a).

**Figure 1:**
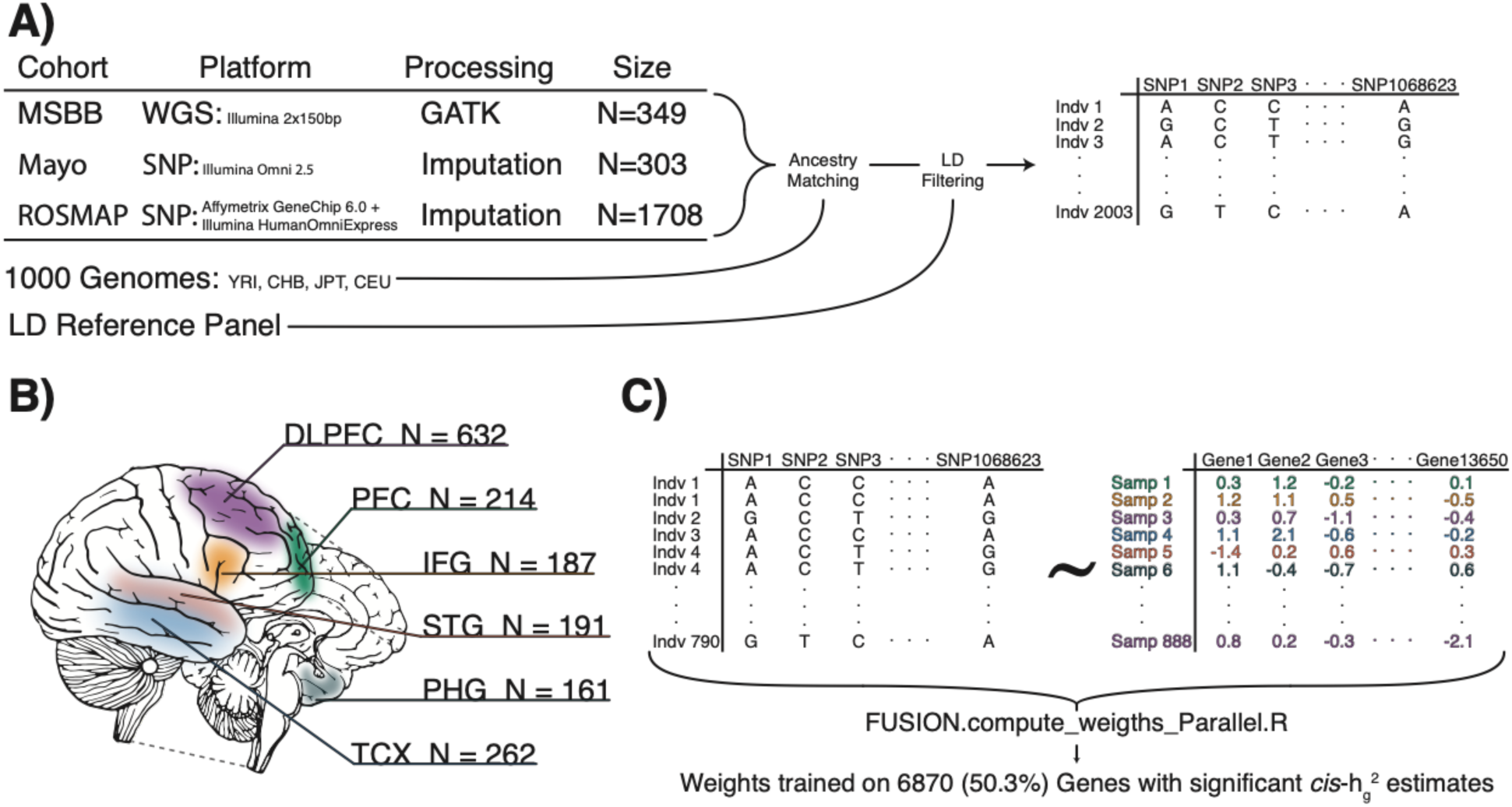
Experimental Design **A)** Sample cohorts from MSBB, Mayo, and ROSMAP were combined across platforms by consensus SNP variant sites. Ancestry analysis was performed and sites within the 2003 CEU ancestrally matched populations were filtered for consensus with the LD reference panel. **B)** RNA-Seq samples originated from 6 distinct Neocortical regions. **C)** The training set data for training TWAS weights consisted of 888 RNA-Seq samples matched to 789 individual variant profiles.

Expression profiling from high-throughput sequencing from all three studies; ROSMAP (DLPFC N=632), Mayo (TCX N=262), and MSBB (PHG N=161, IFG N=187, STG N=191, FP N=214) were normalized within study. For description of the individual expression data and processing see supplementary methods sections 3-4. Iterative normalization was performed to regress significant covariates for individual studies including, but not limited to, diagnosis, age at death, sex, and post mortem interval (Table S1). Post quality control and normalization, 888 RNA-Seq samples were matched to 790 individuals (Fig. 1c, Table S4). Genotypes were represented multiple times for a subset of MSBB individuals to create the 888 unique genotype-expression pairs. Among the 60 ancestrally matched MSBB genotypes; 15 were profiled in all four tissues, 22 were profiled in three tissues, 10 were profiles in 2 tissues, and 13 were only profiled in one of the four tissues.

### Training TWAS Weights

Weights predicting gene expression were trained on matched genotype-RNA-Seq profiles and then used to impute the expression components of all 2003 individuals in our CEU ancestral cohort The training set consisted of 790 ancestrally clustered genotypes matched to 888 normalized, scaled RNA-Seq profiles with diagnosis regressed. The FUSION software[10] was modified to accommodate the presence of multiple RNA-Seq profiles across different regions for the same individual by ensuring that all samples from a given individual were present within a single cross-validation fold during training and model optimization. The FUSION pipeline script was altered both cross-validating cohorts of multiple-samples per-individual, this capacity also ensured that each cross-fold validation sample was within 5% of the size every other fold and could accept pre-scaled expression values with specification of an additional flag *--scale 1*. Gemma (v0.98.1) calculated the *cis*-heritability of scaled expression using all SNPs denoted in the LD-Reference panel and within 1MB window centered on the gene’s TSS for all 13650 autosomal genes. Weights were trained using all five TWAS models (blup, bslmm, lasso, top1, enet) for the 6780 genes with a *cis*-heritability estimate greater than 1%. To support the computational requirements of all five models the FUSION software was altered to run on 5 threads and run in 14x parallel across an AWS c5.18xlarge (72 core 144MB) EC-2 instance. All supporting files for training weights are available on Synapse[15] as well as trained weights in RData files which can be used to impute expression components with a user provided genotype profile.

### Expression Imputation and TWAS Gene Associations

The heritable component of gene expression was imputed for 6780 genes with trained weights. Despite being trained on only the 790 individuals with 888 matched RNA-Seq profiles, expression components based on cis-genoype were able to be imputed for the entire combined Mayo, MSBB, and ROSMAP genotype cohort of 2003 CEU-ancestry matched induviduals on an AWS r3.8xlarge (32 core 144MB) EC-2 instance. Association of AD cases versus control using Kunkle et.al GWAS summary statistics[6] was performed with the FUSION.assoc_test.R script[10]. AD case and control designation was specified with strict neuropathological diagnosis criteria cutoffs as specified in supplementary table 5. Only 635 out of 2003 ancestrally matched individuals from the combined Mayo, MSBB, and ROSMAP genotypes were designated as AD cases (N=404) or controls (N=231).

### CMC DLPFC Validation

CMC count data was ingested and processed similar to the AMP-AD transcriptome data. An iterative normalization model was deployed to identify significant covariates and regress them from the expression data before scaling the data (Table S2). Genotype data was profiled with Affymetrix GeneChip Mapping 5.0K Array and a custom version of the Illumina Infinium CoreExome-24 v1.1 BeadChip (#WG-331-1111). Raw data was filtered to remove SNPs with: zero alternate alleles, MAF <1%, genotyping rate < 0.95, Hardy-Weinberg p-value < 1 x 10-6, and individuals with genotyping rate < 0.95. Imputation was performed using eagle, Minimac and the HRC Reference Panel[16]. Imputed variant data was filtered for SNPs present in the LD reference panel using Plink (v1.9). Expression values were imputed, and kendall correlation values were calculated comparing imputed gene expression to the scaled, assayed expression values. Correlation test values were FDR corrected for the number of matched comparisons N=6643.

### JCP and SMR Analysis

To assess the independence of these associations within their respective 1MB windows JCP testing was performed[15,17] In order to replicate our AD associations, SMR[15,18] was run on all 6780 genes with weights and analyzed for TWAS preliminary hits. Correction for 18 multiple comparisons was applied for replication of associated genes(Table S2). JCP analysis was run on all candidate hit regions with FUSION.post_process.R as previously described[10,17].

### GWAS Enrichments

In order to examine whether the subset of genes with trained TWAS weights were enriched with variants more likely to regulate gene expression within their centered 1MB window compared to brain expressed genes without trained weights or the rest of the genome, summary statistics for both Kunkle *et. al[6]* and Styrkarsdottir *et. al[6,19]* were partitioned into 3 groups. SNPs that were within 1MB of the TSS of genes which had trained TWAS weights, SNPs that were within 1MB of the TSS of Autosomal genes which did not have trained TWAS weights, and autosomal SNPs that weren’t within 1MB of an expressed gene (Intergenic). Wilcoxon Rank-Sum tests were performed comparing p-value distributions between SNPs near genes with trained weights versus those without and intergenic.

### Gene Set Expansion and Cell-Type Analysis

To examine functional enrichments and identify potential processes driving AD, we used coexpression to build out a gene set focused on each TWAS associated gene. We first wanted to consider the possibility that another gene within the 1MB window was co-regulated with our identified gene of interest, and therefore could be the causative gene. For all 6 tissues, Variable Bayes Spike Regression was used to calculate the bootstrapped partial regression of each TWAS associated gene to all other genes. If the mean correlation a gene within the 1MB cis-regulatory region was greater than 0.1, the additional gene was also correlated to all other transcribed genes for all six tissues. The only gene that met this criteria and was added in further analysis was APOE, within the 1MB window surrounding APOC1. For each gene or in the case of the APOC1 locus, APOC1 and APOE, the mean partial correlations across all 6 neocortical tissues were used to enrich for co-expressed functionally related genes. The elbow plots indicating the decay of included genes as the standard deviation of mean partial correlations moves away from the centered mean of zero. This was used to draw cutoffs for inclusion into the expanded gene set at values of 0.7 (TREM2), 1.3 (KNOP1, CD2AP, MTCH2, EED), and 1.7 (APOC1) (Fig. 3a).

**Figure 2:**
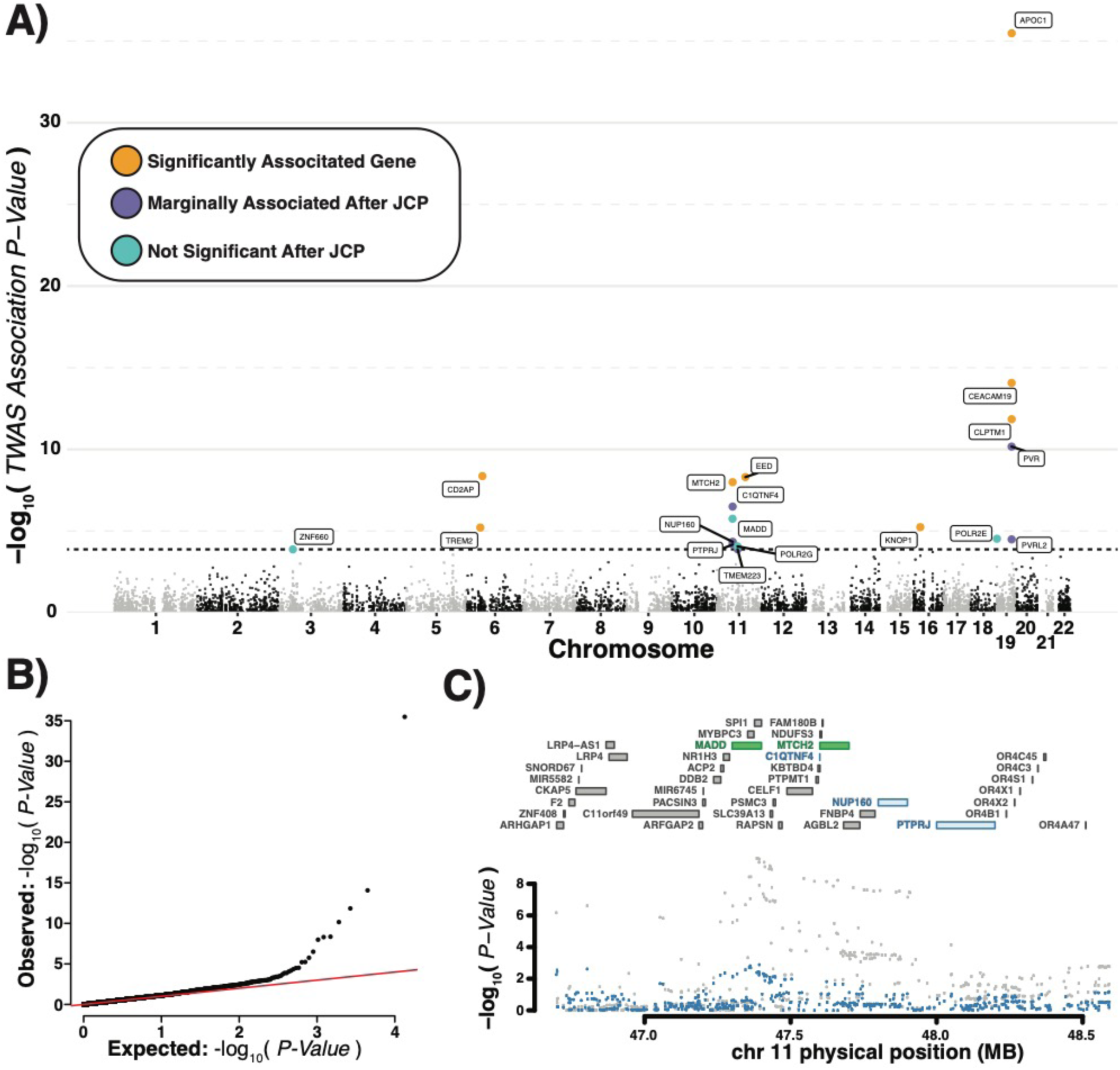
Transcriptome-wide association study results **A)** Log 10 TWAS association p-values by gene shown by genomic location are indicated in black and grey. Features passing initial correction for multiple comparisons (above the dotted line), but marginally significant after Joint Conditional Probability (JCP) are shown in purple. Those features which are no longer significant after JCP are shown in light blue, while genes surviving JCP are shown in Yellow. **B)** QQplot of all TWAS p-values. **C)** An example plot of a region tested for JCP. The candidate genes found to be marginally significant, NUP160 and PTPRJ, are colored blue while those found to be jointly significant, MADD and MTCH2, are colored green (*Upper*), while individual snp p-values are colored grey before and blue after conditioning (*Lower*).

**Figure 3:**
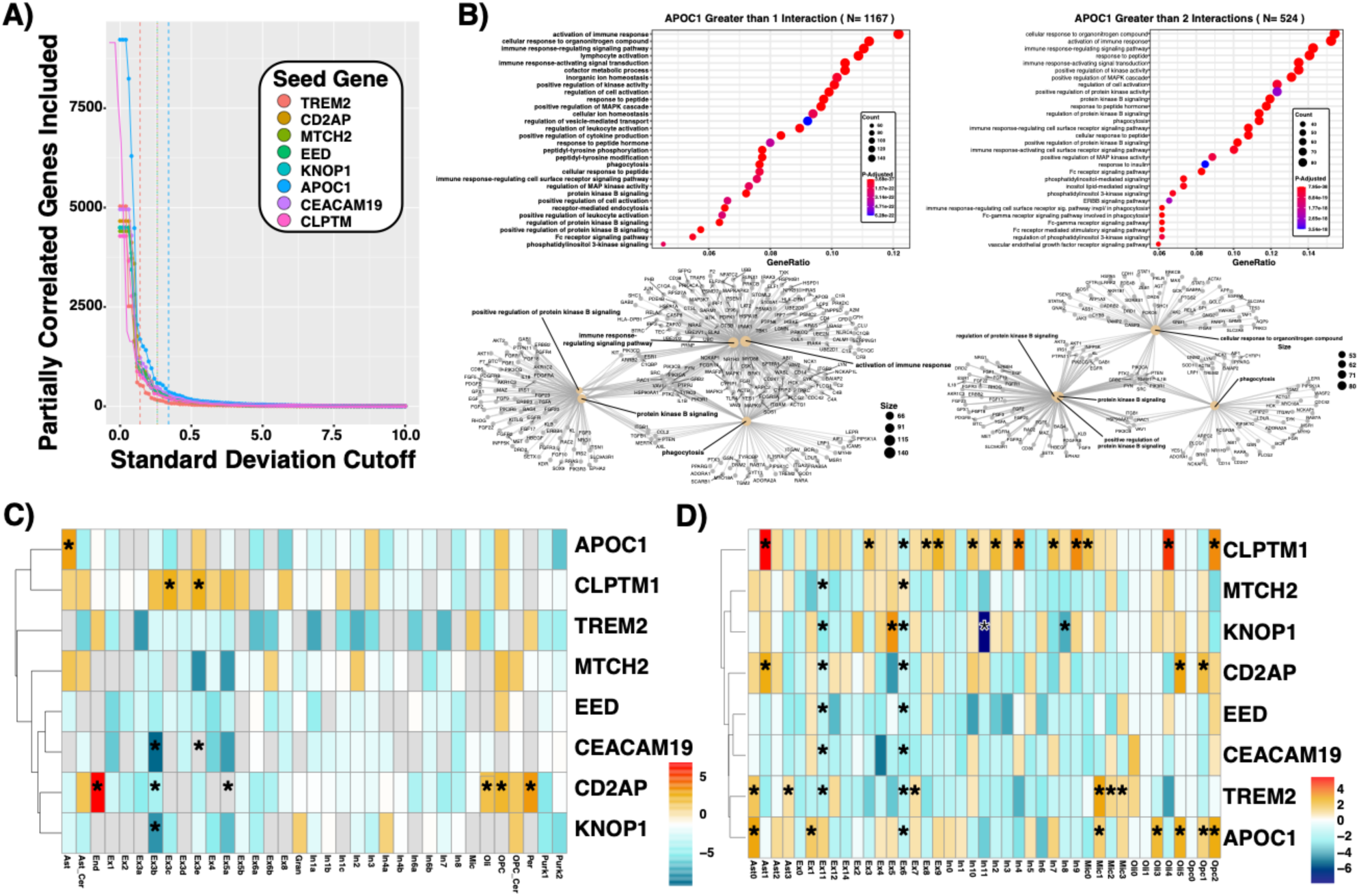
Cell-specific and cell process enrichment analysis. **A)** Gene-set cutoffs of partial correlation to associated gene as a function of standard deviations away from the mean partial correlation. These are the cutoffs used for cell type specific enrichments seen in C and D. **B)** EnrichR cell process enrichments of APOC1 expanded gene set of the 50 highest partially correlated genes and then expanded to all protein interaction partners from the pathway commons database which are represented more than once (Left) and twice (Right). **C)** Cell type specific enrichments of expanded gene sets using Lake *et. al* cell type specific marker gene sets. Grey denotes zero overlapping genes between gene set and cell type-marker gene set significance in a grey square infers depletion from the expected overlap by chance. **D)** Cell type specific enrichments of expanded gene sets using Mathys *et. al* cell type specific marker gene sets.

For cell-type enrichments, TWAS expanded co-expression gene sets were analyzed for enrichment in cluster-specific marker genes from Lake *et. al[20]* and Mathys *et. al[21]*. Odds ratios were calculated with the percent overlap of a gene set and cell-type specific marker gene list divided by the expected percentage of overlap. Significance was calculated with a two sided exact binomial test, FDR < 0.05 corrected for 35 (Lake) and 41 (Mathys) comparisons.

For cell process enrichments a multiscale gene set expansion was employed, foundationally based on small scale enrichment of expanding the candidate TWAS gene to the top 50 bootstrapped partial regression inclusion statistics averaged across all tissues. The second expansion was performed by leveraging pathway commons protein-protein interaction databases from Pathway Commons[22]. All pairwise gene-gene interactions containing a gene member with the initial gene set expansion were extracted and the total gene set was expanded into a broad and narrow range gene set by including any gene which appears more than once (broad) or more than twice (narrow). Ensembl gene IDs were translated to gene symbols and filtered for brain relevance by a filter requiring them to be expressed in at least one brain region. These gene sets were then submitted to EnrichR[23,24] to find cell process enrichments with the background set being any gene set to the list of all genes expressed in any one of the 6 brain regions analyzed.

## Results

### Training and Validating TWAS Weights

While a total 6818 (49.95%) genes had *cis*-heritability estimates greater than 1% and therefore had weights trained for them, only 6780 of 13650 (49.67%) could have expression components with non-zero variance imputed into either the CEU ancestrally matched genotype cohort of 2003 individuals or the training set of 888 profiles. Imputed expression components based solely on based on *cis*-genotype contribution were correlated to actual expression values for all 888 individuals in our training set. Kendall correlations were calculated for all 6780 imputed genes to actual gene expression, after a an FDR < 0.05 correction for multiple comparisons accounting for initial complete set of transcripts (N=13650), 6775 (99.93%) were significantly correlated with the actual expression values at an FDR < 0.05 and 6716 (99.06%) were significantly correlated after bonferroni correction (Fig. S2a). The distribution of correlations was right-skewed towards one (X= 0.24±0.09) (Fig. S2b). Comparing imputed expression components to actual gene expression for four representative weights (Fig. S4) confirmed that weights were not biased by tissue type or cohort, and our regression normalization and expression scaling coupled with changes to the FUSION trained weights specific to the continuous heritable expression difference across our training set (Fig. S3[a-d]).

CMC DLPFC data was used to validate imputed expression components. The validation data set comprised 515 individually matched genotype to expression profiles with a set of 6756 expressed genes overlapping the trained weights, representing 99.6% of all trained weights. Kendall Rank Correlation values between imputed expression components and observed expression was right-skewed towards one (X= 0.13±0.11) (Fig. S4b). FDR correction for multiple comparisons yielded 4874 (72.14%) genes with significant, positive correlations between imputed and actual DLPFC expression (Fig. S4a). Genes with a significant p-value (FDR < 0.05) and positive correlation values all had correlation values greater than 0.061 (Fig. S4c).

Given the threshold of expression heritability required to train weights for a given gene, it could be expected that variants regulating gene expression would be enriched within 1MB of genes with trained weights versus genes which did not meet the heritability threshold. We looked for enrichment of low p-value SNPs within the 1MB widow centered on genes with trained weights to test this assumption. Variants from Kunkle *et. al[6]* were binned into three groups(Fig. S5a). The first were the autosomal variants within the 1MB window of genes which had trained weights, the second consisted of autosomal variants within the 1MB window of genes which did not meet the 1% heritability threshold to have weights trained for them. The final group of variants was termed intergenic and consisted of all variants outside of a 1MB window centered on any of the 13650 expressed genes in the training dataset regardless of whether predictive weights were able to be trained for the gene. Variants within 1Mb of genes meeting the heritability threshold versus were significantly enriched in lower GWAS P-Values than those within 1MB of genes below the heritability threshold (p < 2.22^-16^ Wilcoxon Rank-Sum) as well as intergenic variants (p < 3.70^-15^ Wilcoxon Rank-Sum). To confirm this result, the same analysis was performed with variants from Styrkarsdottir *et. al*’s[19] bone density GWAS analysis(Fig. S5b). This outgroup confirmed enrichment for associated variants within genes of higher heritability versus those of lower heritability (p < 2.22^-16^ Wilcoxon Rank-Sum) as well as intergenic variants (p < 2.22^-16^ Wilcoxon Rank-Sum). Despite potential edge cases, such as genes regulated from long distance LD, genes in MHC regions, and *trans*-regulatory effects; this analysis suggests that our weights are enriched for genes under a higher degree of *cis*-genetic modular control, although the rate at which linkage-disequilibrium affects SNP independence is unknown. The outgroup bone density GWAS dataset confirms that this enrichment is for genetic variants controlling gene expression irrespective of AD phenotype, and tissue context, supporting these weights as a general resource across neocortical regions for generalizable use across multiple phenotypes.

### Alzheimer’s Disease TWAS

Imputed gene expression components were associated with AD through implementation of the FUSION pipeline. This analysis yielded 18 preliminarily significant associations across 8 regions after correction for multiple comparisons (FDR < 0.05) (Fig. 2a, Table 1)[15]. JCP testing resulted in dropping 6 preliminary associations as a result of marginal association (Fig. 2c, Fig. S6). FDR correction of the JCP p-value resulted in dropping an additional 4 targets due to non-significance. Association of all remaining genes was further supported by SMR[15] (Table S3). The remaining 8 genes comprising 6 distinct non-overlapping 1MB genomic regions and are significantly associated to AD after JCP and SMR with FDR corrected p-values are: APOC1 (JCP=2.22e-22, SMR=3.41e-4), EED (JCP=3.373e-5, SMR=2.50e-4), CD2AP (JCP=2.96e-5, SMR=2.66e-4), CEACAM19 (JCP=3.27e-5, SMR=1.00e-2), CLPTM1 (JCP=4.04e-3, SMR=2.58e-3), MTCH2 (JCP=0.011, SMR=3.32e-6), TREM2 (JCP=0.021, SMR=2.64e-3), KNOP1(JCP=0.039, SMR=2.50e-4). All gene associations except MTCH2 and KNOP1 replicated[15] and survived JCP[15] testing when summary statistics from Jansen *et. al.* were used instead of Kunkle *et. al.* summary statistics.

**Table 1:** Heritability (*h*^*2*^), Best performing model, before and after JCP Z-Values and p-values for all initially significant AD associated genes. Blue denotes those only marginally significant after JCP, while Green represents independently significant genes.

| ID | Pass JCP | $h^2$ | MODEL | TWAS.Z | TWAS.P | FDR TWAS.P | COND BETA | COND BETA.SE | COND.Z | COND.P | FDR COND.P |
| --- | --- | --- | --- | --- | --- | --- | --- | --- | --- | --- | --- |
| APOC1 | YES | 0.0613 | bslmm | 12.56 | 3.34<sup>-36</sup> | 2.25<sup>-32</sup> | 12.6 | 1 | 12.6 | 3.30<sup>-36</sup> | 2.22<sup>-32</sup> |
| EED | YES | 0.0234 | bslmm | -5.85 | 4.99<sup>-09</sup> | 5.60<sup>-06</sup> | -5.8 | 1 | -5.8 | 5.00<sup>-09</sup> | 3.37<sup>-05</sup> |
| CD2AP | YES | 0.0372 | bslmm | 5.87 | 4.35<sup>-09</sup> | 5.60<sup>-06</sup> | 5.9 | 1 | 5.9 | 4.40<sup>-09</sup> | 2.96<sup>-05</sup> |
| CEACAM19 | YES | 0.1585 | bslmm | 7.76 | 8.52<sup>-15</sup> | 2.87<sup>-11</sup> | 6.1 | 1.1 | 5.7 | 9.70<sup>-09</sup> | 3.27<sup>-05</sup> |
| CLPTM1 | YES | 0.0355 | bslmm | -7.08 | 1.42<sup>-12</sup> | 3.19<sup>-09</sup> | -5.1 | 1.1 | -4.8 | 1.80<sup>-06</sup> | 4.04<sup>-03</sup> |
| MTCH2 | YES | 0.0390 | bslmm | -5.73 | 1.02<sup>-08</sup> | 9.81<sup>-06</sup> | -4.8 | 1 | -4.7 | 3.30<sup>-06</sup> | 0.011 |
| TREM2 | YES | 0.0795 | bslmm | -4.52 | 6.19<sup>-06</sup> | 3.79<sup>-03</sup> | -4.5 | 1 | -4.5 | 6.20<sup>-06</sup> | 0.021 |
| KNOP1 | YES | 0.3668 | blup | -4.53 | 5.80<sup>-06</sup> | 3.79<sup>-03</sup> | -4.5 | 1 | -4.5 | 5.80<sup>-06</sup> | 0.039 |
| POLR2E | YES | 0.2162 | blup | 4.18 | 2.97<sup>-05</sup> | 0.017 | 4.2 | 1 | 4.2 | 3.00<sup>-05</sup> | 0.051 |
| POLR2G | YES | 0.0448 | bslmm | -3.92 | 8.69<sup>-05</sup> | 0.037 | -3.9 | 1 | -3.9 | 8.70<sup>-05</sup> | 0.195 |
| ZNF660 | YES | 0.1090 | bslmm | -3.82 | 1.33<sup>-04</sup> | 0.049 | -3.8 | 1 | -3.8 | 1.30<sup>-04</sup> | 0.876 |
| MADD | YES | 0.4328 | blup | -4.78 | 1.77<sup>-06</sup> | 1.32<sup>-03</sup> | -3.5 | 1 | -3.4 | 6.40<sup>-04</sup> | 1 |
| PVRL2 | NO | 0.0278 | bslmm | -4.16 | 3.25<sup>-05</sup> | 0.0168 | -5.6 | 0.97 | -5.7 | 9.60<sup>-09</sup> | 6.47<sup>-05</sup> |
| PVR | NO | 0.1082 | blup | -6.53 | 6.76<sup>-11</sup> | 1.14<sup>-07</sup> | -2.2 | 0.85 | -2.6 | 8.80<sup>-03</sup> | 1 |
| TMEM223 | NO | 0.0516 | bslmm | -3.82 | 1.31<sup>-04</sup> | 0.0498 | -2.087 | 0.9 | -2.328 | 0.02 | 1 |
| NUP160 | NO | 0.0362 | bslmm | 4.08 | 4.59<sup>-05</sup> | 0.0220 | 0.849 | 0.72 | 1.179 | 0.24 | 1 |
| PTPRJ | NO | 0.0518 | enet | 3.99 | 6.68<sup>-05</sup> | 0.0300 | 0.254 | 0.6 | 0.426 | 0.67 | 1 |
| C1QTNF4 | NO | 0.0140 | enet | 5.11 | 3.19<sup>-07</sup> | 2.69<sup>-04</sup> | 0.046 | 0.56 | 0.083 | 0.93 | 1 |

### Cell Type Specificity and Pathway Enrichment

Expanded gene sets of genes co-expressed to TWAS nominated AD associated genes were built empirically for each region’s gene set based on membership decay given increasing standard deviation cutoff (Fig. 3a, *methods section: Gene Set Expansion and Cell-Type Analysis*). Distinct cell type enrichments were seen comparing gene sets to the cell-type specific marker lists observed in Lake *et. al[20]* (Fig 3c) and Mathys *et. al* [21](Fig 3d). APOC1 and TREM2 coexpression sets were enriched in astrocyte and microglial markers respectively. The CD2AP gene set was enriched within endothelial, pericytes, and oligodendrocyte expression signatures. KNOP1, CEACAM19, MTCH2, and EED co-expression gene set was not enriched within any of the cell-type specific expression sets, but showed sporadic enrichments in neuronal cell types (Fig. 3c-d). CLPTM1 was enriched across neuronal populations, oligodendrocytes, and astrocytes (Fig 3c-d). Notably, the largest two single-cell marker gene sets, derived from excitatory neuronal populations, Ex6 and Ex11, were more prone to enrichment. This could have been due to their containing significantly more genes than the other gene sets, which contain 850 (Ex6) and 747 (Ex11).

Cell process enriched gene sets were built with a multiscale approach combining pairwise inclusion statistics with protein-protein interactions. Both a wide inclusion cutoff and a more stringent inclusion cutoff produced a permissive and conservative gene set for each candidate gene (see methods [*Gene Set Expansion and Cell-Type Analysis*]). APOC1 was enriched for multiple immune-response signaling pathways, phagocytosis and immune activation consistently across both enriched gene sets (Fig 3b). CD2AP was enriched for cellular responses to lipids, protein localization and responses to multiple molecular compounds (Fig S7). EED, CLPTM1, and CEACAM19 were consistently enriched for RNA, mRNA processing, RNA splicing, RNA-translation processes. In addition to high amounts of transcription relevant overlap between the three candidate genes distinction of enrichments of; viral gene expression response (EED), protein catabolic processes for (CLPTM1), and myeloid/megakaryocyte differentiation (CEACAM19) were observed (Fig S8,S12-13). MTCH2 was enriched purine, deoxy and ribo-deoxynucleotide metabolism (Fig S9). KNOP1 was the smallest gene set within both the permissive and conservative cutoff groups. Nominal enrichments for mitotic cell phase transition and Wnt signaling point to potential a role in cell-cycle progression (Fig S10). TREM2 expanded gene sets showed enrichment for immune response activation, T-Cell and leukocyte activation, as well as cell motility and phagocytosis (Fig S11).

## Discussion

We trained predictive models to impute gene-expression components attributable to *cis*-variation across multiple neocortical tissues. This is the first pan-cortical analysis and is broadly abstractable throughout the neocortex, providing a valuable resource to investigate a multitude neurological conditions and disorders. By including a range of neocortical structures relevant to AD we sought to specifically identify drivers of AD capable of working across these diverse regions. Beyond AD, there are a number of neuropsychiatric conditions; Schizophrenia, depression, and ASD to name a few, which affect the neocortex as a whole. As AD disease status was regressed from training set expression data, our trained weights represent a valuable resource capable of giving insight into the mechanisms of neocortical phenotypes.

We leveraged these weights to perform a TWAS between Alzheimer’s cases versus controls, revealing 8 candidate genes across 6 distinct regions which passed multiple filters for significance after correction for JCP and SMR replication. We used the Jansen *et. al* 2019 AD GWAS study[7] to replicate our findings, and we confirmed six of the eight genes with this data set. The two genes that failed to replicate significant associations, after correction for multiple comparisons, were MTCH2 and KNOP1, which were not identified in Jansen *et. al[7]*, indicating that our methodology is consistent with the input GWAS statistics. Importantly, as imputed expression is dependent on genotype, gene expression is associated with AD directly through underlying regulatory *cis*-genetic risk factors. These methods can have difficulty in training expression weights for relevant genes which have a high variance or are regulated under trans-regulation such as miRNA mediated transcript decay. Larger sample sizes and a wider array of neocortical tissue types may help in mitigating these difficulties, however the vital nature of these biospecimens make it understandably difficult to address completely. Co-regulation within the *cis*-genetic window is a possible confounder in any TWAS analysis, as a more stably expressed co-regulated gene could possibly produce a more robust association than the true causative disease-linked gene[25]. With the exception of APOE being highly coexpressed with APOC1, no other gene within a window is appreciably coexpressed with the TWAS candidate gene, granting confidence in our 8 candidate genes. The inclusion of a diverse set of brain regions into the training set may disrupt co-regulation based on tissue-specific expression and differential disease impact across brain regions could introduce variance into the model. This is a particular strength of our study however as AD pathology affects all of our included regions, therefore fundamental driving risk genes could be expected to be identified across our neocortical tissue-set. While it is important to consider the whole locus in the context of our TWAS associations, this evidence supports our associated genes.

### CD2AP, Chr6

The CD2AP locus contains multiple G-protein coupled receptor (GPR) genes, in addition to CD2AP itself (Fig. S6). JCP analysis supports CD2AP as the most likely linked gene within this locus, an observation that is consistent with broader biological investigations implicating upregulation of CD2AP in AD. All four of the largest GWAS studies performed looking at AD genetic associations have found variants that point to CD2AP[6,7,26,27]. The biological role of CD2AP involves dendritic targeting of APP to the intraluminal vesicles (ILV), which functions as the post-synaptic lysosomal complex, for degradation[28]. Targeting APP to the ILV leads to proteolytic clearance, and decreases the shared time spent with BACE within the endosomal complex, and results in a net decrease in secreted amyloid. Concordantly, knockdown of CD2AP function impairs the targeted degradation of APP, and allows APP and BACE to co-exist within the early endosomal compartment, promoting increases in amyloid production[28]. Accordingly, studies over-expressing CD2AP drive a shift in APP localization from Rab5+ early endosome to Rab7+ late endosome, leading to lysosomal degradation, and decreases amyloid secretion[29]. Autosomal dominant AD mutations, associated with the early onset form of AD, resulted in enlargement of the early endosomal compartment and elevated levels of the BACE cleaved APP carboxy-terminal fragment (CTFbeta) in cortical neurons derived from IPSCs[30]. This work supports the emerging viewpoint that perturbations in endocytosis play a fundamental role in AD biology.

Towards its role in AD, CD2AP operates in concert with BIN1 as a functional regulation mechanism in concert with BIN1. While CD2AP promotes trafficking towards ILV for degradation in the dendrites, BIN1 targets BACE from the early endosome back to the cell surface within axons, preventing the colocalization of APP and BACE in endosomal compartments and decreasing the levels of secreted amyloid. While it may be too narrow a perspective to look only at amyloid biogenesis for linkage with disease mechanism, it at least provides one plausible framework for consideration. Expanded gene set enrichment of CD2AP identifies a range of processes implicated in both the regulation of tissue development as well as responses to lipid and organic cyclic compounds (Fig. S7). Other potential biological roles can be seen in mice, where CD2AP is implicated in blood brain barrier function[31], and in Drosophila where the CD2AP ortholog *Cindr* is implicated in synaptic plasticity and Tau linked neurodegeneration[32,33]. The role of CD2AP in AD biology has been predominantly examined in neurons, while our cell-type enrichments point to the primary involvement of endothelial, oligodendrocyte, pericytes, and astrocytes. Future studies exploring the function of CD2AP in non-neuronal cells may prove useful in developing a broader perspective of CD2AP function in AD pathogenesis.[6,27]

### EED, Chr11

The EED gene was identified by Kunkle *et. al*, but it was not by three other major AD GWAS from the last few years[6,7,26,27]. One possible explanation is that the EED locus contains PICALM, a known AD risk factor, and could lead EED to be overlooked by other types of studies. We do not believe that PICALM explains the TWAS risk identified here for a number of reasons. PICALM is positively associated with AD as it is involved in clathrin-mediated endocytosis of APP, and subsequent generation of amyloid[34,35], however EED’s TWAS association Z-value is -5.85 (Table 1). This infers that overexpression of the loci’s regulated gene target is protective against AD and this means the valence of the effect runs in the wrong direction for a PICALM association. Alternatively, EED is a component of the polycomb repressive complex 2 (PRC2) that functions as a histone methylase depositing the repressive mark H3K27[34–36]. Targeting EED and the PRC2 complex plays a role in synaptic plasticity, as genetic ablation of EED or MII (another element of the PRC2 complex) directly impacts long-term potentiation, a surrogate measure to hippocampal memory function[37]. Additionally, EED and the PRC2 complex promotes neurogenesis within the hippocampus, potentially making the brain more resistant to age related neurodegenerative changes[38,39]. Interestingly, the EED expanded gene set was enriched for many processes involving translation and RNA splicing, two biological domains that would be impacted by heterochromatin regulation (Fig. S8). This locus remains interesting given the potential roles of both EED and PICALM in AD biology, and further study is needed to fully understand the roles of each gene’s contribution to the disease risk.

### MTCH2, Chr11

MTCH2, or mitochondrial carrier 2, is a SLC25 family member of transporters, which localizes to the inner mitochondrial membrane. MTCH2 has been identified as an AD risk factor[6,26]. Previous studies implicated risk variants regulating SPI1[40] and CELF1 expression, with fine mapping potentially implicating CELF1[41] from the MTCH2 locus in AD. While siRNA knockdown of SPI1 in microglia have implicated it as potential target for treatment in AD, MTCH2 is negatively associated with AD (Z-Value = -5.73 Table 1), meaning that reduced expression of MTCH2 contributes to AD pathology, reinforcing our interpretation that MTCH2 is the associated gene. The biological role of MTCH2 in the brain is unclear. MTCH2 is known to contribute to adipocyte function, regulation of lipid metabolism[40,42], and to be genetically associated with obesity[43]. However, MTCH2 clearly has a role outside of adipocytes, as inhibition of MTCH2 increases products of metabolism, such as pyruvate and pyruvate dehydrogenase[44] in both brain and muscle. Our gene set enrichment analysis further supports MTCH2 involvement across a wide array of metabolic processes (Fig. S9). It is not tremendously unexpected that MTCH2 has a direct impact on neurological function as the brain is one of the most metabolically active organs in the body. MTCH2 knockout mice exhibited deficits in both metabolic processes and hippocampal dependent spatial learning tasks[42,45,46]. There are known links between nutrition, specifically cholesterol consumption levels, in AD[47], which can also present health risks for cardiovascular function, another well-known risk factor for AD. Concordantly, there are associations observed between obesity[48] and AD, suggesting that MTCH2 variants associated with AD and obesity may be acting, at least in part, through a common mechanism, in which modulation of MTCH2 expression impacts AD risk by driving changes in BMI[49]. This does not account for the spatial learning deficits, suggesting that this is not the whole answer. Yet, consistent with a non-neuronal locus of effect, we did observe that the MTCH2 expanded gene set is enriched for microglial and oligodendrocyte cell-type markers (Fig. 3d). MTCH2 knockout mice exhibit elevated levels of microglia and diminished synaptic density in the basal forebrain, both of which could be explained by perturbations in microglial and oligodendrocyte function[46]. The mechanisms through which MTCH2 exerts its influence upon AD pathogenesis are currently not fully elucidated, additional studies will be necessary to fully understand the relevant biology.

### KNOP1, Chr16

KNOP1 is a lysine rich nucleolar protein lacking direct publications and while currently little is known of its biological function, down regulation is associated with AD risk (Table 1). The KNOP1 gene resides within the IQCK locus, and Kunkel *et. al[6]* found linkage between KNOP1 and AD; however, none of the other recent GWAS studies found KNOP1 associated with AD. IQCK was a novel genome-wide locus from the Kunkle *et. al[6]* genetic meta-analysis. What data exists for KNOP1 suggests that its associates condensed chromatin during mitosis, which is partially supported by AP2M1 a KNOP1 yeast two-hybrid binding partner, Human Reference Interactome (HuRI)[6,50] search binds with a large number of H2B associated proteins[50,51]. This preliminary evidence aligns with the KNOP1 expanded gene set enrichment which shows a strong signal for mitotic cell phase transition and regulation of DNA-binding transcription-factor activity (Fig. S10). Intriguingly, the entire locus, similar to variants in the MTCH2 locus, is implicated in obesity[52,53]. We hope that the finding that KNOP1 is associated with the AD risk will inspire future studies into its specific function.

### TREM2, Chr6

Identification of TREM2 in this study is consistent with previous work and knowledge in AD, as TREM2 is one of the most widely studied genes in Alzheimer’s disease, with links to both amyloid and tau pathology. TREM2 is expressed almost exclusively in microglia and it is a sentinel gene linking neuroinflammation to AD[54]. TREM2 null mutant mice crossed onto APP-PS1 AD transgenics exhibit deficits in microglial recruitment to amyloid plaques, and increased spread of pathological tau[55]. Coding variants in TREM2 are estimated to confer a 2-4 fold increase in AD risk, higher than any gene other than APOE[56,57]. Interestingly, APOE binds to TREM2, leading to activation and recruitment of the microglial cells, which elicits both phagocytic and proinflammatory responses[54]. TREM2 also binds to amyloid directly, with nanomolar affinity, and activates microglial clearance of amyloid deposition[55,58]. Consistent with TREM2 function, our expanded TREM2 gene set expression was enriched for immune myeloid cellular lineages and cell process enrichments of immune activation and leukocyte migration (Fig. 3[c-d], Fig. S11). Interestingly our TREM2 Z-score is -4.52 (Table 1), which appears to contradict previous work. However, TWAS analysis associates only the genetic component of TREM2 expression, inferring that TREM2 genetic risk could function through disruption and dysregulation of TREM2 endogenous function. This is consistent with the disruptive TREM2 coding variants and recessive loss of function associations with AD.

### APOC1 - CEACAM19 - CLPTM1, Chr19

APOC1, CLPTM1 and CEACAM19 were identified within this study, and all three genes reside within 600 Kb, less than a 1 Mb distance threshold, of each other. Despite this proximity, there appears to be complicated co-regulation in this region with upregulation of APOC1 and CEACAM19 associated with AD but downregulation of CLPTM1 associated with AD. Interesting, all three of these genes were also identified in recent GWAS studies[7,26]. This locus is particularly infamous for harboring the APOE genes, the largest known LOAD risk allele. APOE was the most coexpressed gene to APOC1, and this co-expressed pair had the highest average coexpression of any gene within a *cis*-locus to an associated gene, strongly supporting the co-regulation of APOC1 and APOE. Little is known about the specific role of APOC1 in AD; however, as it is also a lipid carrier transport protein that, like APOE, is known to recruit the innate immune system, it may also have a role in regulation of microglial activation. More specifically, there are isolated studies which demonstrate linkage between APOC1 and declining cognition and memory in specific ethnic groups[59,60]. Consistent with the APOC1 role in lipid transport and immune activation, our expanded cell process enrichments specifically identified phagocytosis and activation of immune response (Fig. 3B), supporting a shared role between APOE and APOC1 as the biological contributor to enhanced AD risk, and making the disentanglement of each gene’s unique AD risk contribution that much more difficult.

While APOC1, CEACAM19, and CLPTM1 are associated with AD, both in this study and previous studies[60–62], a causal association to AD remains unclear. However, there is a small literature pointing to a role for CLPTM1 in the regulation of GABA receptor trafficking from the ER to the plasma membrane, suggesting that CLPTM1 could regulate inhibitory neurotransmission [63,64]. The regulation of GABA currents could be a synaptic scaling factor, adjusting the responsiveness of synaptic firing. Ge *et. al*[63] found that increasing CLPTM1 levels decreased miniature inhibitory postsynaptic potentials, while reciprocally, decreasing CLPTM1 levels elevated GABA currents in the post-synaptic neuron, strongly suggesting that CLPTM1 negatively regulates GABAergic signaling. A recent literature meta-analysis looking at neurotransmitter synaptic dysregulation in AD, found decreased levels GABA in AD patients, supporting the potential dysregulation of GABAergic signaling in AD[65]

Far less is known about CEACAM19, and any biological connection to AD. What work has been done is predominantly focused upon oncology. However, it was observed in oncogenic investigations that CEACAM19 does interact with the AKT/PI3K pathway[13], a pathway known to play a regulatory role in tau phosphorylation. While these studies certainly do not prove a connection between CLPTM1 or CEACAM19 and AD, they do suggest a plausible biological connection that warrants further investigation. Unlike other loci, the APOC1/CLPTM1/CEACAM19 locus has an extremely tight cluster of associated genetic risk variants clustered near the APOE/TOMM40/APOC1 gene loci. Consequently, it is difficult for TWAS studies to separate out linkage disequilibrium and potential coregulation from variant risk exerted through specific causal genes. We do not believe this invalidates our findings, yet it is a difficulty for all TWAS studies examining this loci.

## Conclusions

We have presented here a TWAS analysis of Alzheimer’s using weights trained from RNA-Seq expression values derived from 6 distinct cortical regions to associate genetic risk to expression differences in 404 cases and 231 controls. This methodology has shown its power in resolving additional mechanistic insights in the impact of risk variants on transcripts responsible for AD pathology. We provide a resource of trained expression weights for 6818 genes which is broadly abstractable across the neocortex and when used in combination with summary GWAS statistics can perform powerful associations across a broad range of neocortical phenotypes.

## Abbreviations

AD: Alzheimer’s disease
CEU: Central European
FDR: False Discovery Rate
LOAD: Late Onset Alzheimer’s Disease
EOAD: Early-Onset Alzheimer’s Disease
GWAS: Genome Wide Association Studies
TWAS: Transcriptome Wide Association Study
DLPFC: Dorsolateral Prefrontal Cortex
TCX: Temporal Cortex
PFC: Prefrontal Cortex
STG: Superior Temporal Cortex
IFG: Inferior Temporal Gyrus
PHG: Parahippocampal Gyrus
AMP-AD: Accelerating Medicines Partnership - Alzheimer’s Disease
CMC: CommonMind Consortium
JCP: Joint Probability Correlation
SMR: Summary Mendelian Randomization
GPR: G-protein Coupled Receptor
ILV: Intraluminal Vesicles

## Declarations

### Ethics approval and consent to participate

Not applicable.

### Consent for publication

Not applicable.

### Availability of data and materials

Weights and analysis files are available via the AD Knowledge Portal (https://adknowledgeportal.synapse.org). The AD Knowledge Portal is a platform for accessing data, analyses, and tools generated by the Accelerating Medicines Partnership (AMP-AD) Target Discovery Program and other National Institute on Aging (NIA)-supported programs to enable open-science practices and accelerate translational learning. The data, analyses and tools are shared early in the research cycle without a publication embargo on secondary use. Data is available for general research use according to the following requirements for data access and data attribution (https://adknowledgeportal.synapse.org/DataAccess/Instructions). See 10.7303/syn22163073 for information about data and data sources https://www.synapse.org/#!Synapse:syn22163073

### Authors’ contributions

JG, BAL, SKS, APW, and TSW designed the study. JG, KSM, WLP generated the data. JG performed the experiments and analyzed the data. JG preformed the statistical analyses and interpreted the data with input from BAL, SKS, TSW, and APW. JG and JCW wrote the manuscript with input from all co-authors. JG and JCW revised the paper with input from all co-authors. All authors read and approved the final manuscript.

## Acknowledgements

First and foremost, the authors would like to acknowledge the patients and their families who provided the invaluable tissue samples to support this work. Rush Alzheimer’s Disease Center, Rush University Medical Center, Chicago provided all ROSMAP data collected, which was funded by; NIA grants: P30AG10161, R01AG15819, R01AG17917, R01AG30146, R01AG36836, U01AG32984, U01AG46152, the Illinois Department of Public Health, and the Translational Genomics Research Institute. Mayo Study data were provided by the following sources: The Mayo Clinic Alzheimer’s Disease Genetic Studies, led by Dr. Nilufer Ertekin Taner and Dr. Steven G. Younkin, Mayo Clinic, Jacksonville, FL using samples from the Mayo Clinic Study of Aging, the Mayo Clinic Alzheimer’s Disease Research Center, and the Mayo Clinic Brain Bank. Mayo Data collection was funded through; NIA grants: P50 AG016574, R01 AG032990, U01 58AG046139, R01 AG018023, U01 AG006576, U01 AG006786, R01 AG025711, R01 AG017216, R01AG003949, NINDS grant R01 NS080820, CurePSP Foundation, and support from Mayo Foundation.

