## Supplemental Materials for "Multi-Tissue Neocortical Transcriptome-Wide Association Study Implicates 8 Genes Across 6 Genomic Loci in Alzheimer’s Disease"

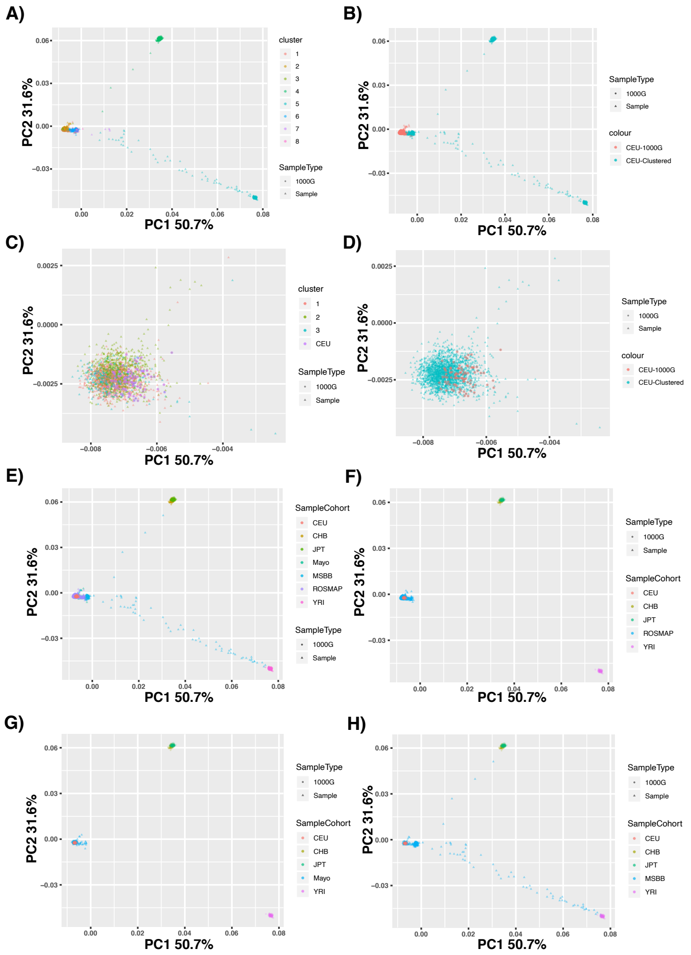

**Figure S1. Ancestral Matching of Combined Cohort**

Ancestral matching of combined sample genome set. **A)** Clustering of combined MSBB, Mayo, ROSMAP, and 1000G (YRI, CEU, CHB, JPT) genotype sets on principle components describing more than 1% of variance with GemTools.R software revealed 8 distinct clusters. **B)** Three of these clusters contained CEU reference samples (soft-red), while the other 5 clusters did not (cyan). **C)** These 3 clusters were not distinct from each-other between the first 2 principal components. **D)** Populations were largely ascertained from CEU ancestral populations. **E)** ROSMAP and **F)** Mayo cohorts were extremely similar in ancestral composition while **H)** MSBB had greater amounts of admixture reflecting the difference in ascertainment.

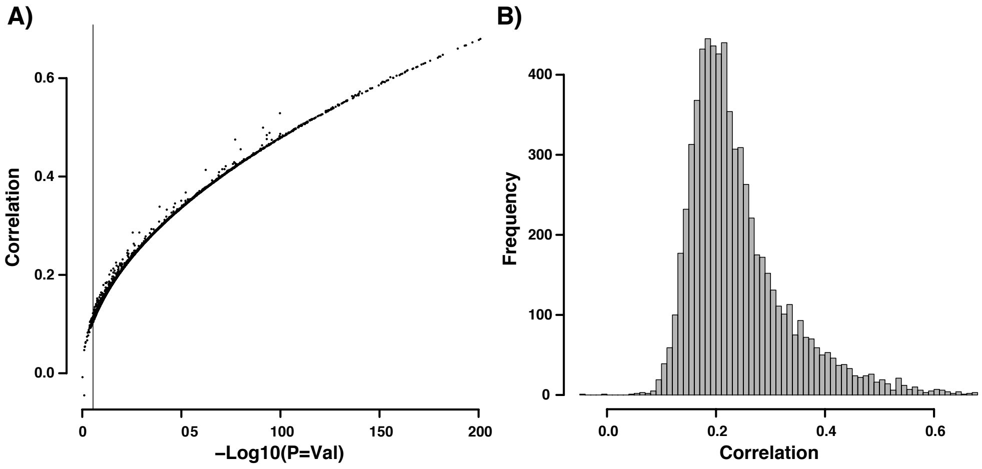

**Figure S2. Actual Expression Versus Imputed Expression**

Correlation of imputed expression components versus actual expression in ROSMAP DLPFC. **A)** Log10 p-values as a function of matched Kendall correlation values for all genes, with bonferroni significance denoted by the vertical line. **B)** Distribution of Kendall correlation values across all imputed genes compared to actual expression values.

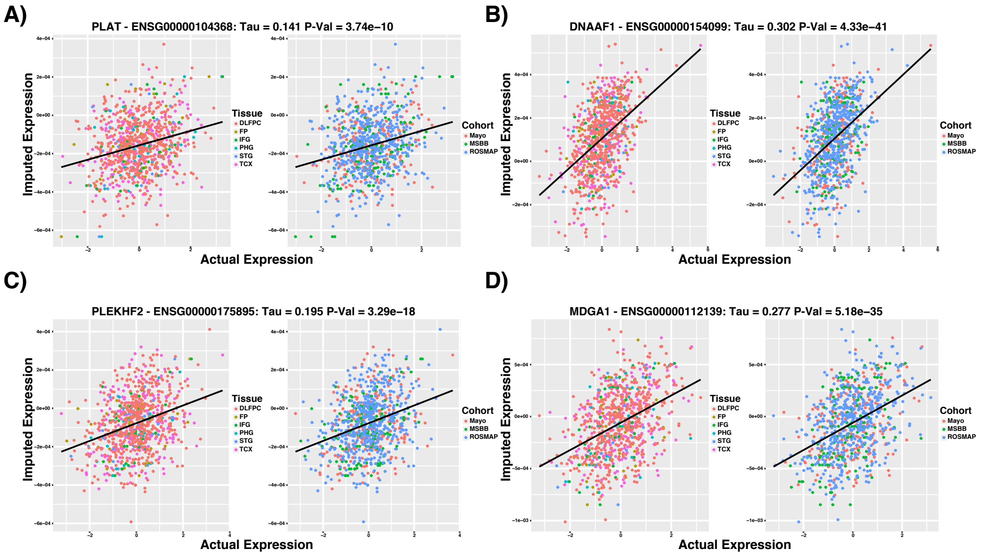

**Figure S3. Validation Actual Versus Imputed Examples**

Imputed versus actual expression for four representative sets of weights. Plot titles include gene names and ENSG names as well as Kendall Correlation Tau values and uncorrected P-Values. Colors for each set of plots A thru D represent the sample tissue of origin *Left* and the individual's cohort of origin *Right*.

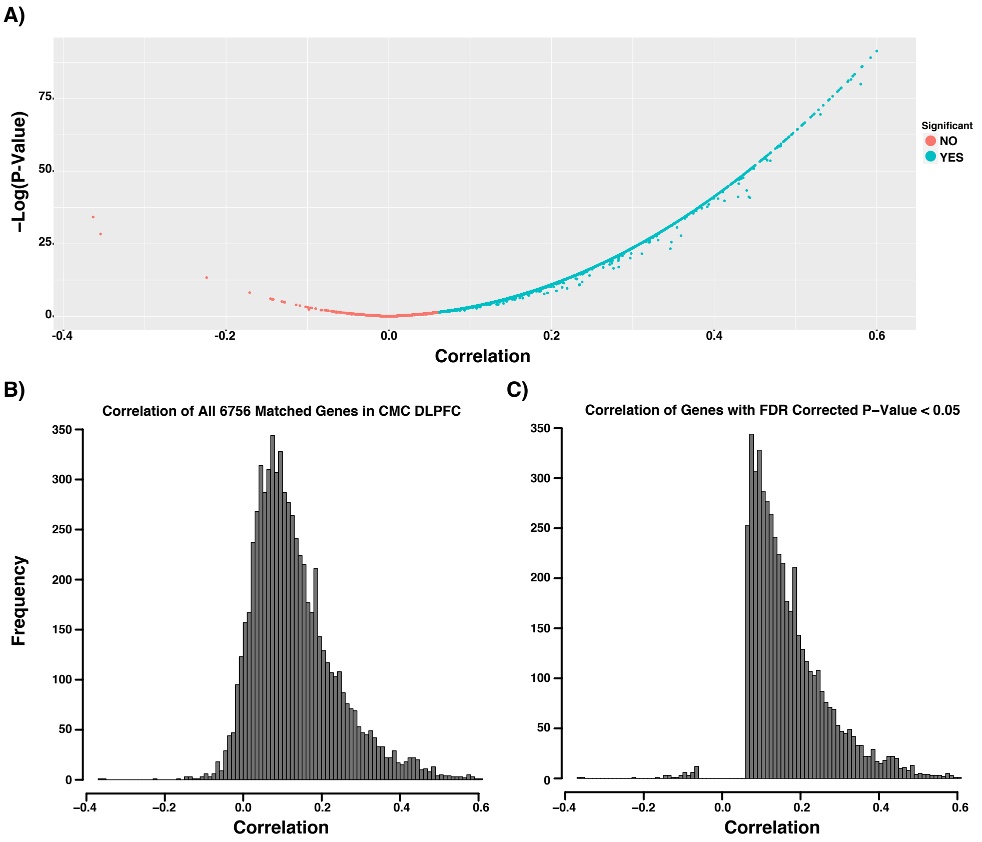

**Figure S4. CMC DLPFC Validation**

Validation of weights in CommonMind Consortium DLPFC RNA-Seq **A)** Log10 p-value as a function of associated kendall correlation value, significant p-values (FDR ¡ 0.05) in cyan, non-significant correlations are in soft-red. **B)** Distribution of all correlation values across paired weights to expressed DLPFC genes. **C)** Distribution of all correlation values across paired weights to expressed DLPFC genes with a significant p-value after correction for multiple comparisons (FDR ¡ 0.05).

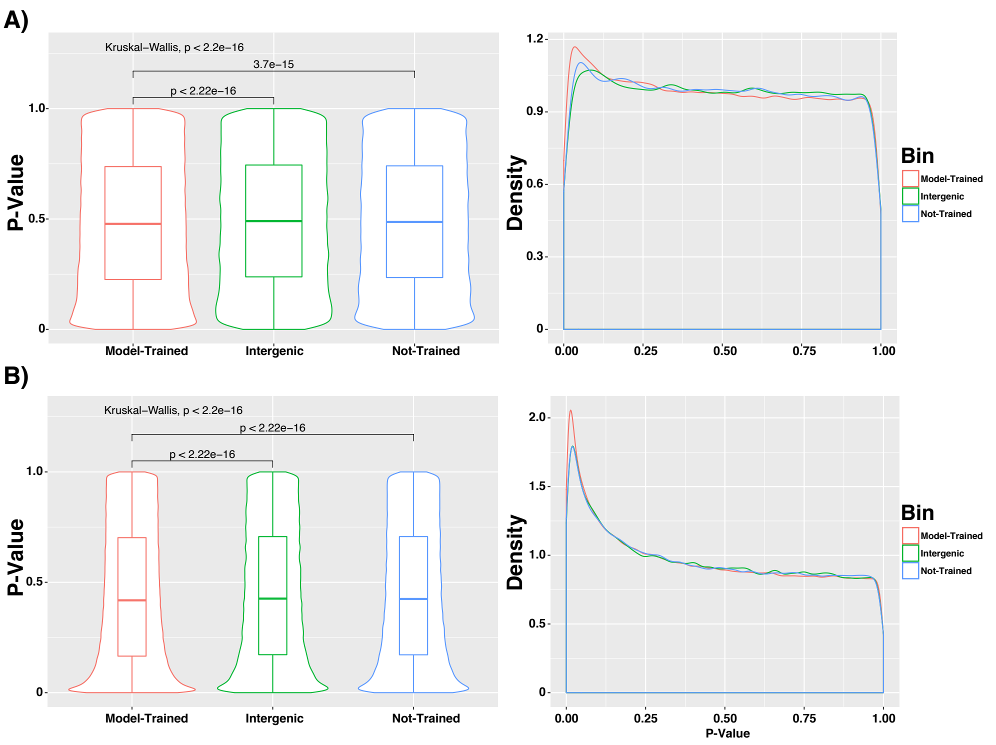

**Figure S5. Enrichment of GWAS SNPs Near Genes With Trained Weights**
GWAS p-values of SNPs within 1MB of genes which had expression-weights trained (Red), genes which failed to have weights trained for them (Blue), and all other SNPs (Green) from Kunkle *et. al* **(A)** and Styrkarsdottir *et. al* **(B)** GWAS studies. Left panels are boxplots with p-values from Wilcox-rank sum tests denoting significance between individual groups and Kruskal-Wallis p-values denoting significance between all groups, right panels show the p-value density distributions.

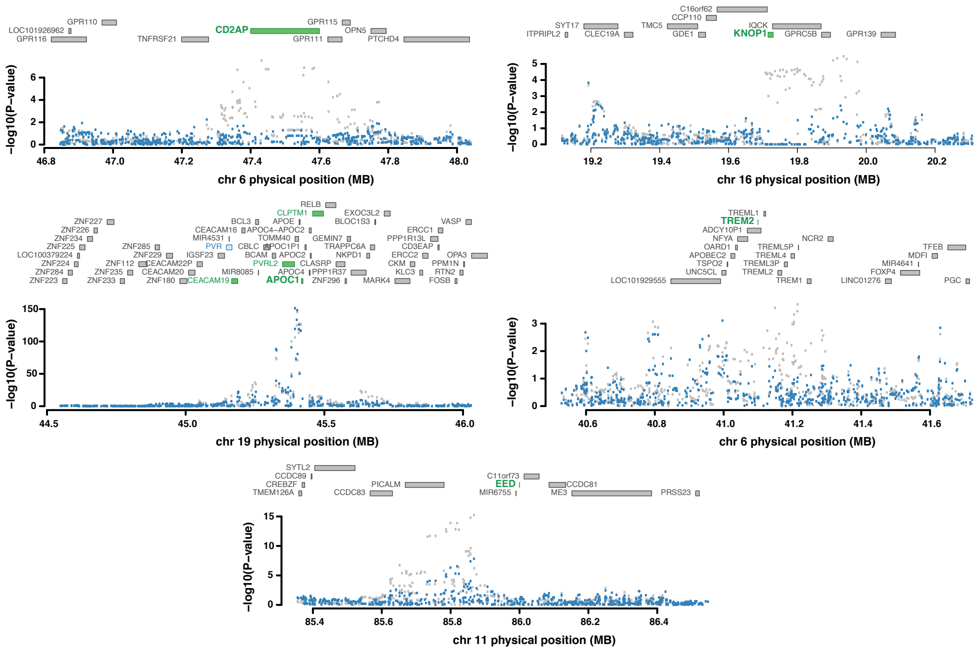

**Figure S6. Joint Conditional Probability Analysis**

Joint Conditional Probability Analysis of the associated genomic region to CD2AP (upper left), KNOP1 (upper right), APOC1, CLPTM1, CEACAM19 (mid-left), TREM2 (mid-right), and EED1 (lower). For each panel of candidate genes, those found to be marginally significant are colored blue, while those found to be jointly significant are colored green *(Upper)*, while individual snp p-values are colored grey before and blue after conditioning *(Lower)*.

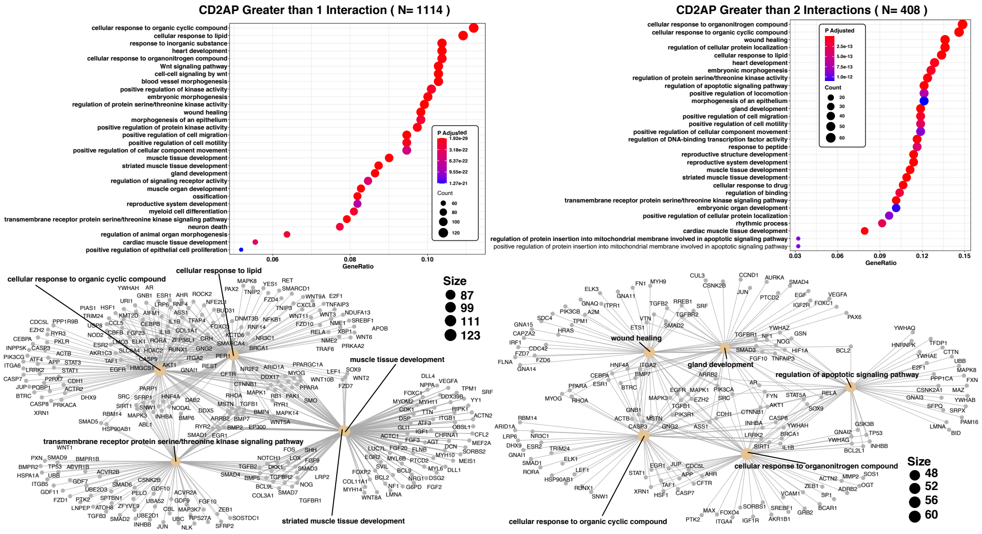

**Figure S7. CD2AP Cell Process Enrichment**

EnrichR cell process enrichments of CD2AP expanded gene set of the 50 highest partially correlated genes and then expanded to all protein interaction partners from the pathway commons database which are represented more than once *(Left)* and twice *(Right)*.

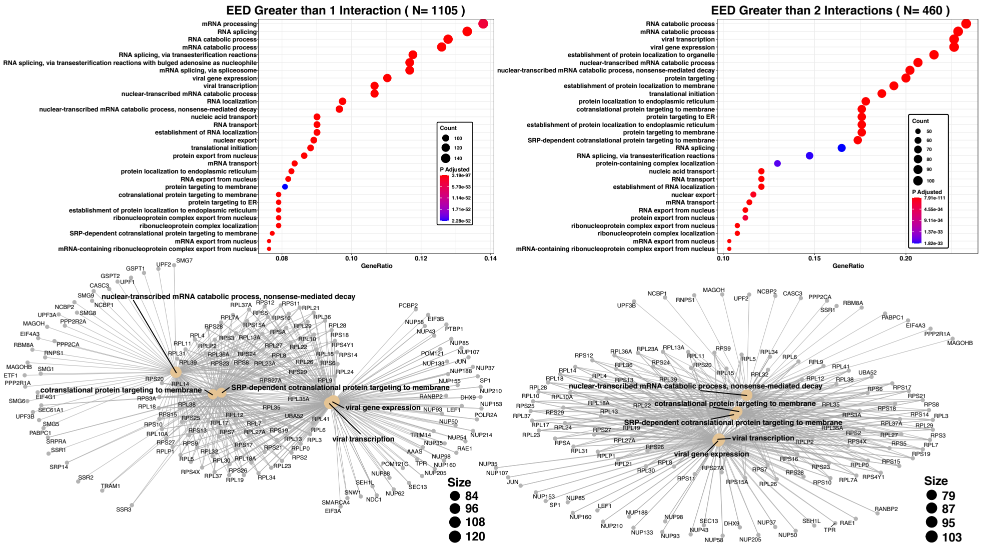

**Figure S8. EED Cell Process Enrichment**

EnrichR cell process enrichments of EED expanded gene set of the 50 highest partially correlated genes and then expanded to all protein interaction partners from the pathway commons database which are represented more than once *(Left)* and twice *(Right)*.

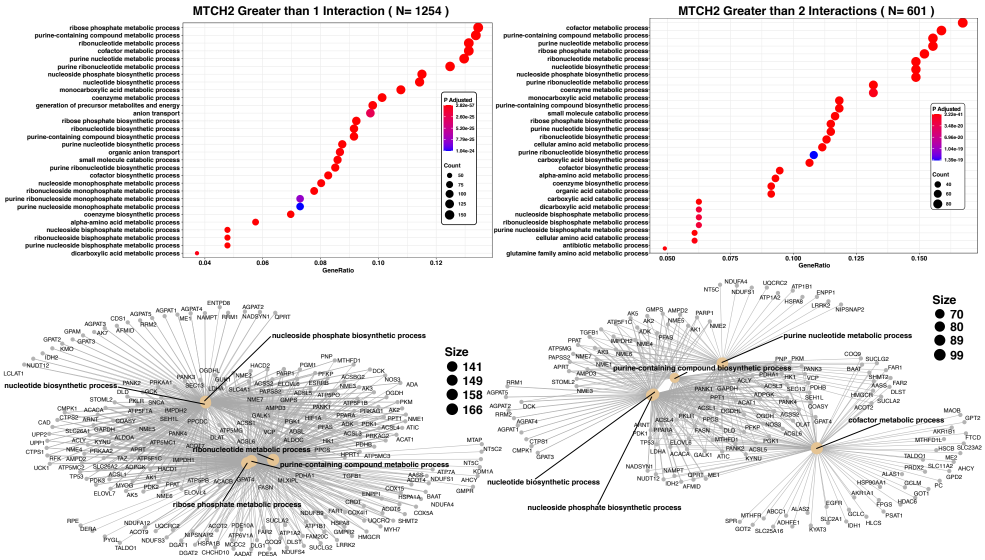

**Figure S9. MTCH2 Cell Process Enrichment**

EnrichR cell process enrichments of MTCH2 expanded gene set of the 50 highest partially correlated genes and then expanded to all protein interaction partners from the pathway commons database which are represented more than once *(Left)* and twice *(Right)*.

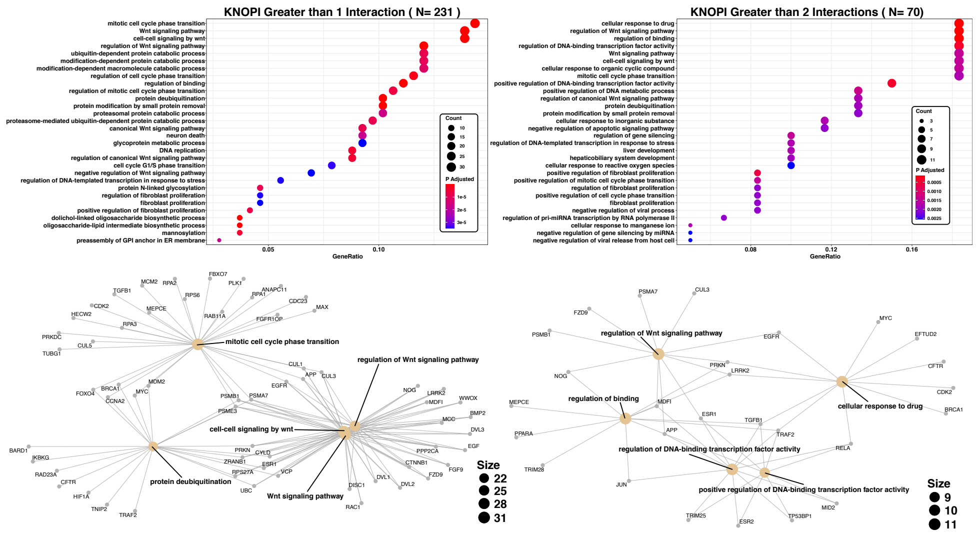

**Figure S10. KNOP1 Cell Process Enrichment**

EnrichR cell process enrichments of KNOP1 expanded gene set of the 50 highest partially correlated genes and then expanded to all protein interaction partners from the pathway commons database which are represented more than once *(Left)* and twice *(Right)*.

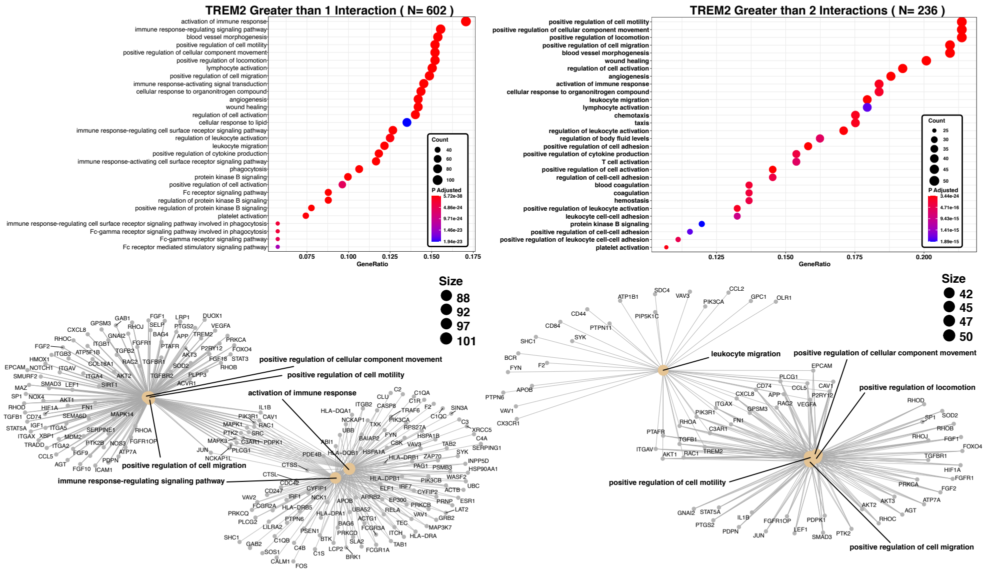

**Figure S11. TREM2 Cell Process Enrichment**

EnrichR cell process enrichments of TREM2 expanded gene set of the 50 highest partially correlated genes and then expanded to all protein interaction partners from the pathway commons database which are represented more than once *(Left)* and twice *(Right)*.

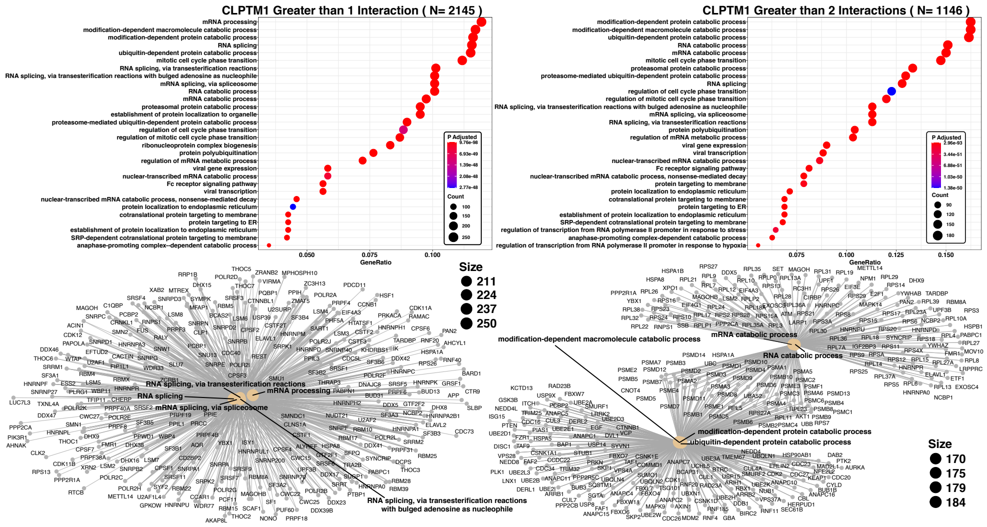

**Figure S12. CLPTM1 Cell Process Enrichment**

EnrichR cell process enrichments of CLPTM1 expanded gene set of the 50 highest partially correlated genes and then expanded to all protein interaction partners from the pathway commons database which are represented more than once *(Left)* and twice *(Right)*.

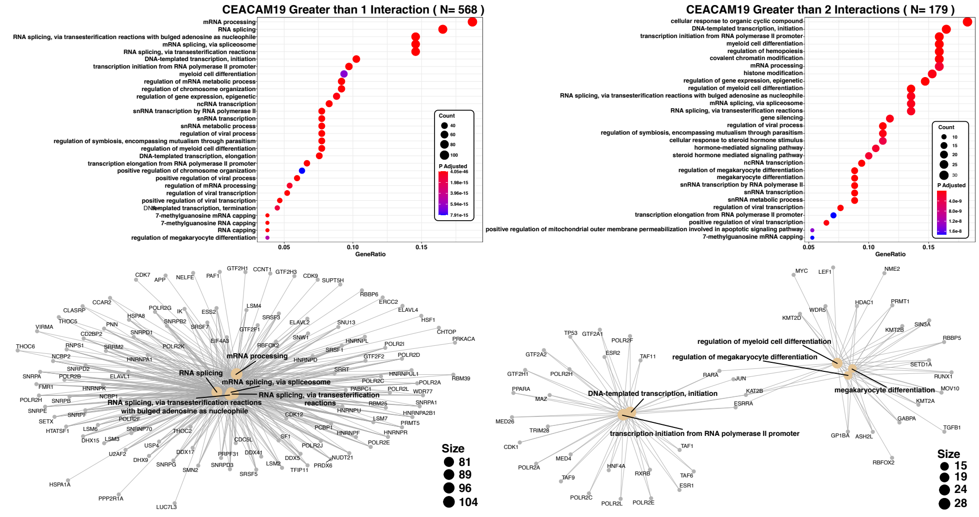

**Figure S13. CEACAM19 Cell Process Enrichment**

EnrichR cell process enrichments of CEACAM19 expanded gene set of the 50 highest partially correlated genes and then expanded to all protein interaction partners from the pathway commons database which are represented more than once *(Left)* and twice *(Right)*.

**Supplemental Methods**

**4 RNA-Seq Processing** **17**

### 1. Genotype Data

#### *1.1. ROSMAP*

ROSMAP genotype data were assayed using the Affymetrix GeneChip 6.0 (n=1709) and Illumina HumanOmniExpress (n=382) genotyping array platforms. Both batches of samples were QC'd as reported in De Jager *et. al*[1] and imputed using BEAGLE software, version 3.3.2.

#### *1.2. Mayo*

Mayo Genotype data were assayed using Illumina's Human Omni 2.5 + Exome array (n=314) samples were QC'd as previously described *(10.7303/syn22163073)* and imputed via the Michigan Imputation Server v1.0.0[2] with the Haplotype Reference Consortium (hrc.r1.1.2016) panel[3].

#### *1.3. MSBB*

MSBB Whole Genome Sequencing for 349 individuals was prepared using the KAPA Hyper PCR-free chemistry targeting an insert size of 450bp fragments. Samples were prepared from 650ng of purified genomic DNA, sonicated with a Covaris LE220 sonicator. Fragments were size selected via AMPureX beads, prepped for Illumina sequencing and quality control was performed with qPCR using the Universal KAPA Library Quantification Kit and the Agilent 2100 BioAnalyzer Fragment Analyzer. Library sequencing was performed on an Illumina HiSeq X sequencer using v2.5 chemistry for paired-end 150bp reads. Data processing was performed with the NYGC automated pipeline. Briefly, reads were aligned to GRCh37 human reference using the Burrows-Wheeler Aligner (BWA-MEM v0.7.08) and variants were called through deployment of GATK best-practices with Picard tools v1.83 and Genome Analysis Toolkit (GATK v3.4.0). Duplicated reads were ignored, local realignment around INDELs and base quality score recalibration (BQSR) were deployed. Specific to variant calling, Genome Analysis Toolkit HaplotypeCaller module was run on each sample in GVCF mode to produce gVCFs, which were combined into a multisample VCF and filtered utilizing Variant Quality Score Recalibration (VQSR) *(10.7303/syn22163073)*.

### 2. Genotpype Processing

MSBB WGS vcf files were converted to plink format, filtered for SNVs, and merged across autosomes utilizing plink v1.9[4]. Snps were filtered for overlap with imputed genotyping data using location and RSIDs were assigned through matching coordinated with custom python scripts. MSBB, imputed ROSMAP, and imputed Mayo variant data were combined and filtered independently for overlap with 1000G data and the FUSION LD Reference Panel[5] for ancestry and training gene weights respectively.

### 3. RNA-Seq Data

#### *3.1. ROSMAP*

All ROSMAP samples were derived from DLPFC (n=637) and prepped according to the Broad Institute's strand specific dUTP method[6]. Briefly, this method enriches for messenger RNA through poly-A selection, synthesizes first strand cDNA and uses dUTP for second strand-specific cDNA and is fragmented, end repaired and adaptor ligated for Illumina sequencing. Out of 637 samples, 582 met the quality control threshold of an RNA integrity (RIN) score ¿5 (Bioanalyzer) and quantity threshold (5ug). Paired-end 101bp sequencing was performed on Illumina HiSeq and minimum sequencing depth was 50 million reads.

#### *3.2. Mayo*

All Mayo samples 275 (CBE) and 276 (TCX) were processed with TruSeq RNA Sample Prep Kit V2 (Illumina, San Diego, CA). Samples enriched for messenger RNA via poly-A selection. Library concentration and fragment size was measured using an Agilent Bioanalyzer DNA 1000 chip. All samples were paired end sequenced on an Illumina HiSeq 4000, read lengths were 101 base pairs.

#### *3.3. MSBB*

RNA from 301 individuals was extracted from FP (n=265), STG (n=264), PHG (n=267), and IFG (n=230) with the TruSeq RNA Sample Preparation Kit v2 (Illumina, San Diego, CA). Coding and long non-coding RNA was enriched for with the Ribo-Zero rRNA Removal Kit (Illumina, San Diego, CA). Random hexamers were used to synthesize cDNA, the library was fragmented and size selected using AMPure XP beads (Beckman Coulter, Brea, CA). Libraries were single-end sequenced on Illumina HiSeq 2500 to a length of 101 base pairs.

### 4. RNA-Seq Processing

RNA-Seq reads were aligned to GRCh38 Gencode v24 with the STAR aligner (v2.5.1b) with the quant mode flag set to GeneCounts to export read counts aligned to gene annotations, and sequencing metrics were calculated with Picard

Tools (v2.2.4). RNA-Seq QC and batch correction was applied by study. Iterative normalization was used to identify and correct for covariates within study as listed in table (Table S1). Of note diagnosis status was regressed out from each cohort to ensure that gene expression heritability estimates were independent of AD diagnosis status. RNA-Seq processing code and reports are available for each cohort *(10.7303/syn22163073)* and the final expression data set contained 13650 genes which were expressed across all tissues. Gene expression values were z-scaled within tissue values in preparation for genotype matching for TWAS weight training.
